# Regional epidemic dynamics and Delta variant diversity resulted in varying rates of spread of Omicron-BA.1 in Mexico

**DOI:** 10.1101/2022.10.18.512746

**Authors:** Selene Zárate, Blanca Taboada, Mauricio Rosales-Rivera, Rodrigo García-López, José Esteban Muñoz-Medina, Alejandro Sanchez-Flores, Alfredo Herrera-Estrella, Bruno Gómez-Gil, Nelly Selem Mojica, Angel Gustavo Salas-Lais, Joel Armando Vazquez-Perez, David Alejandro Cabrera-Gaytán, Larissa Fernandes-Matano, Luis Antonio Uribe-Noguez, Juan Bautista Chale-Dzul, Brenda Irasema Maldonado Meza, Fidencio Mejía-Nepomuceno, Rogelio Pérez-Padilla, Rosa María Gutiérrez-Ríos, Antonio Loza, Susana López, Carlos F. Arias

**Author notes:** **Correspondence**: Selene Zárate, **Correspondence:** Blanca Taboada, **Correspondence:** Carlos F. Arias. **Author Notes**, Selene Zárate and Blanca Taboada contributed equally to this work.

## Abstract

The Omicron subvariant BA.1 of SARS-CoV-2 was first detected in November 2021 and quickly spread worldwide, displacing the Delta variant. In Mexico, this subvariant began spreading during the first week of December 2021 and became dominant in the next three weeks, causing the fourth COVID-19 epidemiological surge in the country. Unlike previous SARS-CoV-2 variants, BA.1 did not acquire local substitutions nor exhibited a geographically distinct circulation pattern in Mexico. However, a regional difference in the speed of the replacement of the Delta variant was observed, as some northern states showed persistence of Delta lineages well into February 2022. Mexican states were divided into four regions (North, Central North, Central South, and Southeast) based on the lineage circulation before the dominance of BA.1 to study possible causes for this difference. For each region, the time to fixation of BA.1, the diversity of Delta sublineages in the weeks preceding BA.1 entry, the population density, and the level of virus circulation during the inter-wave interval were determined. An association between a faster Omicron spread and lower Delta diversity, as well as fewer COVID-19 cases during the Delta-BA.1.x inter-wave period, was observed. For example, the North region exhibited the slowest spread but had the highest diversity of Delta sublineages and the greatest number of inter-wave cases relative to the maximum amount of the virus circulating in the region, whereas the Southeast region showed the opposite. Viral diversity and the relative abundance of the virus in a particular area around the time of the introduction of a new lineage seem to have influenced the spread dynamics. Nonetheless, if there is a significant difference in the fitness of the variants or the time allowed for the competition is sufficient, it seems the fitter virus will eventually become dominant, as observed in the eventual dominance of the BA.1.x variant in Mexico.

**Impact statement:** The surveillance of lineage circulation of SARS-CoV-2 has helped identify variants that have a transmission advantage and are of concern to public health and to track the virus dispersion accurately. However, many factors contributing to differences in lineage spread dynamics beyond the acquisition of specific mutations remain poorly understood. In this work, a description of BA.1 entry and dispersion within Mexico is presented, and which factors potentially affected the spread rates of the Omicron variant BA.1 among geographical regions in the country are analyzed, underlining the importance of population density, the proportion of active cases, and viral lineage diversity and identity before the entry of BA.1.

**Data summary:** This work was carried out using data shared through the GISAID initiative. All sequences and metadate are available through GISAID with the accession EPI_SET_220927gw, accession numbers and metadata are also reported in the supplemental material of this article. Epidemiological data was obtained though the Secretaría de Salud website (https://www.gob.mx/salud/documentos/datos-abiertos-152127),

## Introduction

On November 26, 2021, the World Health Organization declared Omicron (initially designated B.1.1.529 in the Pangolin classification) as the newest variant of concern (VOC) of SARS-CoV-2. This variant was detected a few days prior in South Africa and Botswana and made the headlines for its unprecedented ability to outcompete other circulating variants rapidly [1]. By early December, it had spread to several European countries, causing epidemic surges, and soon became dominant worldwide [2].

As sequences accumulated, phylogenetic analyses showed that the Omicron variant consisted of two main separate clades, designated as BA.1 and BA.2 in the Pangolin nomenclature system. BA.1, labeled clade 21K by Nextclade, was the first of the two to spread and caused epidemic outbreaks throughout the world between December 2021 and January 2022. As of March 30, 2022, the BA.1 lineage had diversified into 54 sublineages (hereafter referred to as BA.1.x). Each one has a distinct mutation pattern and differences in geographical distribution, although no distinctive phenotypic variation has been reported for any of them.

BA.1 is characterized by 44 nonsynonymous substitutions, ten synonymous changes, and seven deletions (ranging between one and three amino acids) compared to the reference strain. This large number of mutations had no precedence in all the described SARS-CoV-2 variants and posed questions regarding its possible origin. Notably, the gene coding for the spike protein (S), which mediates virus cell entry via ACE2 recognition, harbors 30 nonsynonymous substitutions, three deletions, and one insertion. The accumulation of genetic changes in S has been related to increased viral transmission compared with previous VOCs [3]. For instance, in Denmark and Italy, it has been estimated that BA.1 had an effective reproduction number (R_t_) around two to three times greater than the Delta variant [4,5]. The large number of mutations in S resulted in an enhanced antigenic escape, reflected by the high number of infections among the vaccinated population and a sharp decline in the neutralization titers of reported monoclonal antibodies, as well as in the sera of vaccinated subjects [6,7].

Additionally, Omicron BA.1.x transmission advantage seems to be associated with a preference towards a TMPRSS2-independent endosomal entry pathway rather than membrane fusion at the cell surface, which has been associated with a preferential infection of the upper respiratory tract resulting in a reduction in virulence, as described in clinical reports from multiple countries [8–11]. Also, in hamster and murine infection models, Omicron exhibited reduced virulence with respect to previous variants [12,13], and *in vitro* assays demonstrated a limited activation of the NF-κB pathway, which may lead to lower inflammation [14].

By the first week of December, when the Omicron variant began expanding in Mexico, the country had experienced three epidemic surges (peaks). The third outbreak, which occurred during the summer of 2021, was caused by the Delta variant, resulting in the largest number of cases up to that date. However, since vaccination of the most vulnerable population had already begun, in the third surge, the number of hospitalizations and deaths was lower than during the second epidemic peak [15]. Early in December, vaccination efforts reached 60% of the adult population with a complete scheme, and a booster dose campaign had started, aimed at people over 60 years of age in light of the imminent arrival of the Omicron variant.

This study examines the entry and spread of Omicron BA.1.x, which caused the fourth wave of the pandemic in Mexico, describing a geographically homogeneous dispersion of BA.1.x lineages in contrast with earlier waves [16–18]. In addition, it was found that the rate of spread of the BA.1.x lineages varied by region. This variation was associated with the proportion of active cases during the inter-wave period and the diversity of Delta at the time of BA.1 entry.

## Methods

### Epidemiological information

National epidemiological data derived from the governmental SARS-CoV-2 surveillance program in Mexico is made available by the General Epidemiological Directorate (Dirección General de Epidemiología) through the Secretaría de Salud website (https://www.gob.mx/salud/documentos/datos-abiertos-152127), which was accessed on April 24, 2022. Cases confirmed positive to COVID-19 and associated deaths from March 1, 2022, to March 31, 2022, were collated per day for the epidemiological analyses. The information was plotted at national and regional levels using a 7-day rolling average on a sliding window with a step of one day. Epidemiological data processing and graphs were created using R v.4.2.1 [19].

### Sequences data source

All sequences and metadata from the Omicron variant available in the Global Initiative on Sharing All Influenza Data (GISAID) database (accessed on May 18, 2022) were downloaded with a submission dateline on March 31, 2022. In total, the set had 13,631 sequences from Mexico, and 3,191,009 from the rest of the world. Regarding Mexican sequences, 5,752 (42.2%) were generated by the National Institute for Genomic Medicine (INMEGEN), 4,186 (30.1%) by the Mexican Consortium for Genomic Surveillance (CoViGen-Mex), 2,851 (20.9%) by the Institute for Diagnostics and Epidemiological Reference (InDRE), and the remaining (6.8%) by other public and private institutions. The information on the Mexican genomes is included in Supplemental Table S1. To evaluate the displacement of the Delta variant by BA.1.x lineages, sequences and metadata from this variant were also retrieved from GISAID, including sequences from November 2021 to January 2022; in total, 5,510 Mexican sequences of the Delta variant were obtained (Table S2). All sequences and metadata are available through GISAID EPI_SET_220927gw.

### Lineage and mutation frequencies

Lineage clade assignment was carried out using Pangolin PLEARN v1.8 [20] and mutation annotation was done using the NextClade tool v.2.0.0-beta.3 alignment [21]. Variant calling was used to determine the genetic diversity and frequency of BA.1.x sublineages in Mexico and the rest of the world. Regarding mutation analysis, all sequences containing more than 5% ambiguous nucleotides (Ns) were excluded from the analyses, resulting in 12,304 genomes from Mexico.

### Phylogeny and haplotype network

A subset of sequences from Mexico’s three most prevalent BA.1.x sublineages was selected for phylogenetic reconstruction. This subset consists of 1,090 sequences chosen to ensure that every state and month were represented. Additionally, 740 sequences from other countries were included with a distribution based on the relative prevalence of the three sublineages on each continent. Sequences were aligned using MAFFT v.7 with the –addfragments option [22], a maximum-likelihood tree was built using iqtree2 [23] and then was time-scaled with LSD2 [24]. The resulting phylogeny was visualized in R using ggtree [25]. Similarly, a haplotype network was built since haplotypes more accurately reflect epidemiological clustering over short periods of time. Thus, using the same alignment employed for the tree, PopART v1.7 [26] was used to generate a Templeton, Crandall, and Sing (TCS) haplotype network [27].

### Analysis of the regional differences in the introduction of BA.1.x and the displacement of the Delta variant

To characterize the introduction and spread of BA.1.x throughout the country, the relative frequencies of circulating lineages per epidemiological week (W) in each state were considered. For this purpose, only genomes with a collection date in November 2021 (when Delta sublineages were dominant), December 2021 (when BA.1.x began to spread), and January 2022 (when BA.1.x became dominant) were included in the analysis (W44-Nov to W04-Jan). First, weekly maps were created to assess differences in the lineage distribution throughout Mexico. Then, a hierarchical clustering was created using R v.4.1.0, considering lineage diversity and abundance across states. From this result, four geographic regions in the country were determined, each including the following states (Figure S1): i) North (N): Baja California, Baja California Sur, Chihuahua, Coahuila, Durango, Sinaloa, and Sonora; ii) Central North (CN): Aguascalientes, Colima, Guanajuato, Jalisco, Nayarit, Nuevo Leon, San Luis Potosi, Tamaulipas, and Zacatecas; iii) Central South (CS): Guerrero, Hidalgo, Mexico City, Michoacan, Morelos, Oaxaca, Puebla, Queretaro, State of Mexico and Tlaxcala; and iv) Southeast (SE): Campeche, Chiapas, Quintana Roo, Tabasco, Veracruz, and Yucatan.

A Spearman correlation analysis was performed to analyze differences in lineage distribution between regions. Heatmaps were generated to show the abundance of lineages throughout time, using the pandas v.1.3.5, seaborn v.0.11.0, and matplotlib v.3.5.2 libraries of Phyton v.3.7.6. Finally, using the Shannon’s index, the diversity of Delta lineages prior to the entry of BA.1.x in each region was determined. The significance of any differences in the Shannon diversity between regions was evaluated using the *Wilcoxon signed-rank* test and its associated *p*-value with an alpha=0.05.

## Results

### BA.1.x entry into Mexico

According to official records from the Mexican government, the first SARS-CoV-2-positive case was registered in Mexico on February 28, 2020. As of March 31, 2022 (W13-Mar), a cumulative number of 5,659,535 cases and 323,016 deaths had been reported, spanning four epidemic surges (waves) (Figure 1A). The peak average daily cases were 7,275 during the first wave (on July 18, 2020), 16,009 during the second wave (on January 10, 2021), 19,305 during the third (on August 10, 2021), and 61,802 during the fourth wave (on January 15, 2022).

**Figure 1.**
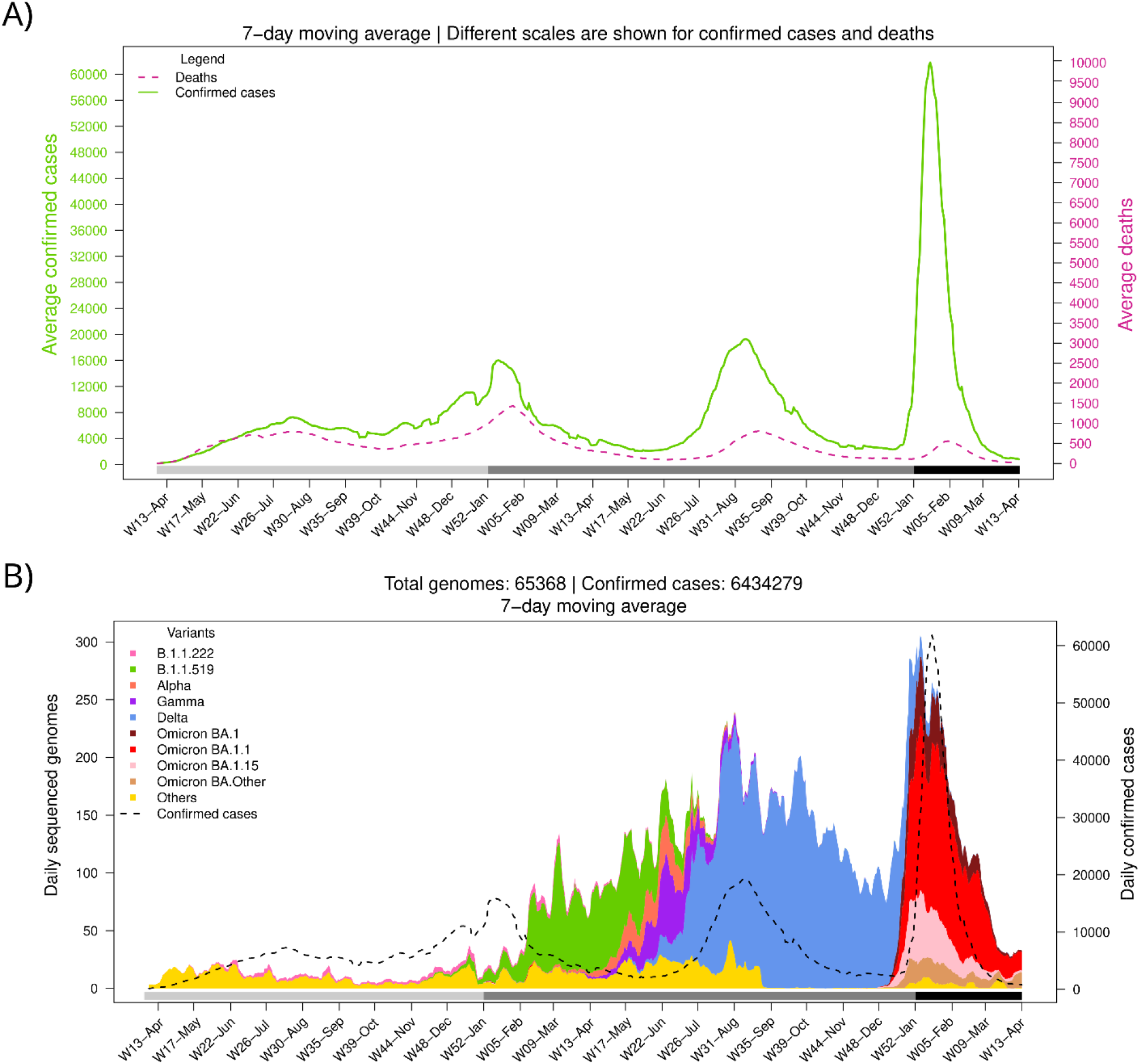
Number of confirmed cases, deaths, and variant diversity of the SARS-CoV-2 virus in Mexico throughout the pandemic. A) Average daily cases and deaths. B) Average daily sequenced genomes are colored by variant. The dashed line shows the total number of confirmed cases. The bottom horizontal bar marks the year (light gray, 2020; gray, 2021; dark gray, 2022).

Different SARS-CoV-2 variants have dominated the Mexican epidemic landscape (Figure 1B). The fourth wave started with the introduction of BA.1.x lineages of the Omicron variant, which displaced Delta (B.1.617.2, and AY.x sublineages), the only variant circulating in the country since the third wave (W25-Jun to W41-Sep 2021). The first confirmed case of BA.1.x was imported on November 16, 2021, during W46-Nov. Omicron’s geographical dominance was swiftly established. During the first week of December (W49-Dec), it was present in 5 out of 32 states (15.6%), spreading to 17 (53.1%) by the second week and to 27 (84.4%) during the third week.

Notably, the fourth wave (W5-Dec2021 to W08-Feb2022) accumulated more cases than the previous waves in a shorter period, with maximum daily records 8.5, 3.9, and 3.2 times greater than those registered during waves 1st, 2nd, and 3rd, respectively. In contrast, reported deaths attributed to COVID-19 have decreased since the second wave, having a case fatality rate of 11.4%, 8.7%, 4.1%, and 1.2% for waves 1st to 4th, respectively. This difference might be due to an increase in the population immunity, in part due to the protection acquired from prior natural SARS-CoV-2 infections and the vaccination campaign, which had covered most adults by the end of the third wave (complete scheme) and included a booster dose for the elderly by late 2021.

### Diversity of BA.1.x in Mexico

Figure S2 illustrates the distribution of the Delta dominant sublineages in Mexico since October 2021 and the fast displacement of this variant by Omicron-BA.1.x sublineages in the weeks following its introduction. During the fourth wave, at least six BA.1.x sublineages circulated in Mexico with a prevalence higher than 1% (Table 1); however, the most abundant three accounted for 91.5% of all sequenced genomes. As shown in Table 1, BA.1.1 accounted for more than 50% of the sequenced genomes, while BA.1 and BA.1.15 had a prevalence of around 17%, each. In contrast, some unique sublineages circulated predominantly in Mexico during the Delta wave compared to the rest of the world [18], these three Omicron sublineages circulated at roughly the same frequencies in the USA. Interestingly, BA.1.1 accounted for around 50% of the sequences in North and South America, except for Canada and Brazil, while it exhibited a low prevalence in the rest of the world. On the other hand, BA.1 was more prevalent in Canada and Brazil (about 38%) than in Mexico and the USA (12% and 16%, respectively).

**Table 1.**
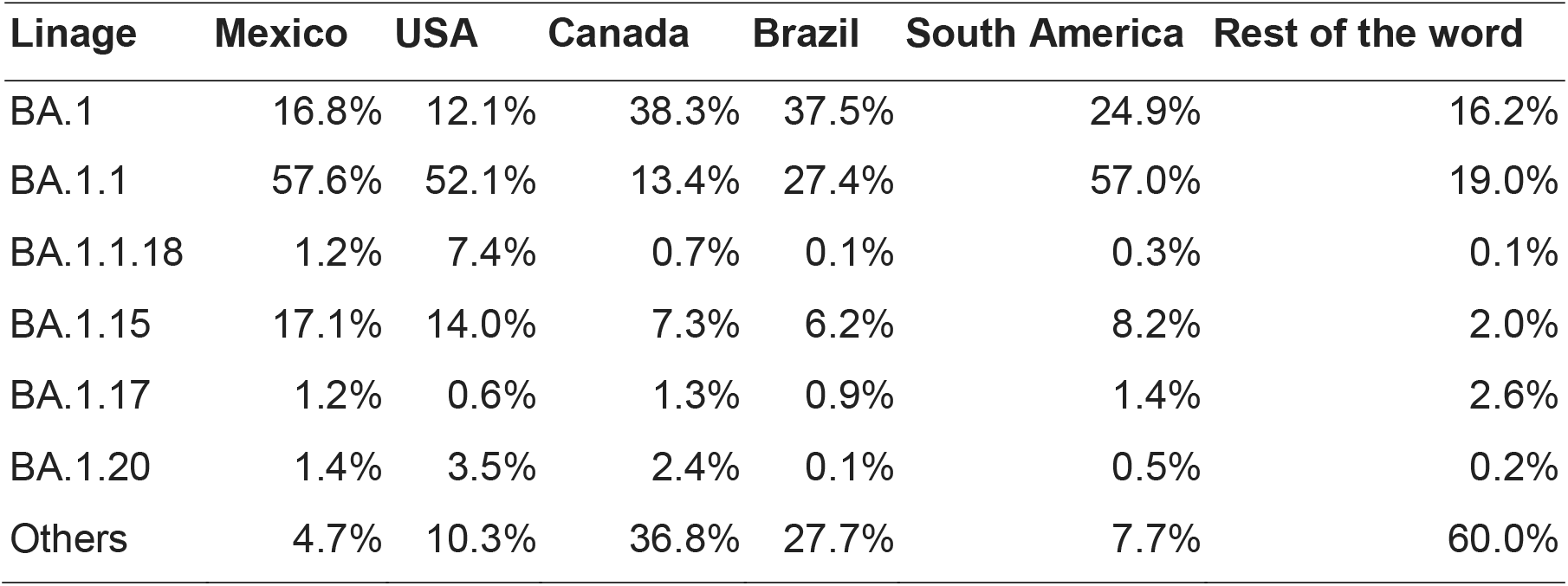
Comparison of the BA.1.x sublineages that circulated in Mexico and other countries. Sublineages with a prevalence smaller than 1% in Mexico were collated into Others.

Whole-genome mutation analysis was carried out using only high-quality genomes (having <3% ambiguous bases) of the six BA.1.x lineages with a prevalence higher than 1% in Mexico (Table 1). We identified 7,144 distinct nucleotide changes, but only 110 (1.5%) were found in more than 1% of the genomes. Regarding nonsynonymous changes, 3,939 were identified, 84 (2.1%) of which had a prevalence higher than 1%. These changes include the 46 amino acid signature substitutions and 7 amino acid deletions (between 1 to 3 aa in length) previously reported in BA.1.x, which were identified in over 90% of the Mexican isolates, as well as a 3-aa insertion that was present in over 40% of sequences (Table S3). Moreover, the additional mutations that define sublineages BA. 1.1 (S:R346K), BA. 1.1.18 (ORF1a:A2554V and S:R346K), BA. 1.15 (ORF3a:L106F and N:D343G), BA.1.17 (ORF1a:V1887I) and BA.1.20 (N:P67S) were also detected proportionally to their prevalence. Only two previously unreported amino acid substitutions were identified in these lineages, with a frequency greater than 5%.

A phylogenetic reconstruction and a haplotype network were built to determine the presence of BA.1.x Mexican haplotypes among the circulating Omicron viruses (Fig S3). To this end, only the three most prevalent sublineages, BA.1, BA.1.1, and BA.1.15, were included in the analysis. Interestingly, these sublineages did not form distinct clusters in the network and were highly polytomic in the phylogeny (Figure S3), probably due to their low genetic diversity. Also, the Mexican sequences were interspersed with those from abroad, similarly pointing to the limited diversification of these lineages. Moreover, when analyzed by geographical region, Mexico showed no difference in sublineage genetic diversity (Figure S3). This limited sequence diversity could be related to the rapid global and national spread of the BA.1.x variant.

### BA.1.x entry in Mexico was asynchronous

A series of maps were built to characterize the dynamics of the BA.1.x spread in Mexico. Figure 2 shows that at epidemiological week 49, all states were dominated by the Delta variant (blue). However, the following week, some states showed a larger proportion of BA.1.x sequences (red). During the last week of December 2021, most states in Central and South Mexico and many in the north had a majority of BA.1.x sequences. By the first week of 2022, the national prevalence of BA.1.x exceeded 50%, and during January, its prevalence continued to rise. Of interest, in some northern states, the replacement of the Delta variant was slower than in others.

**Figure 2.**
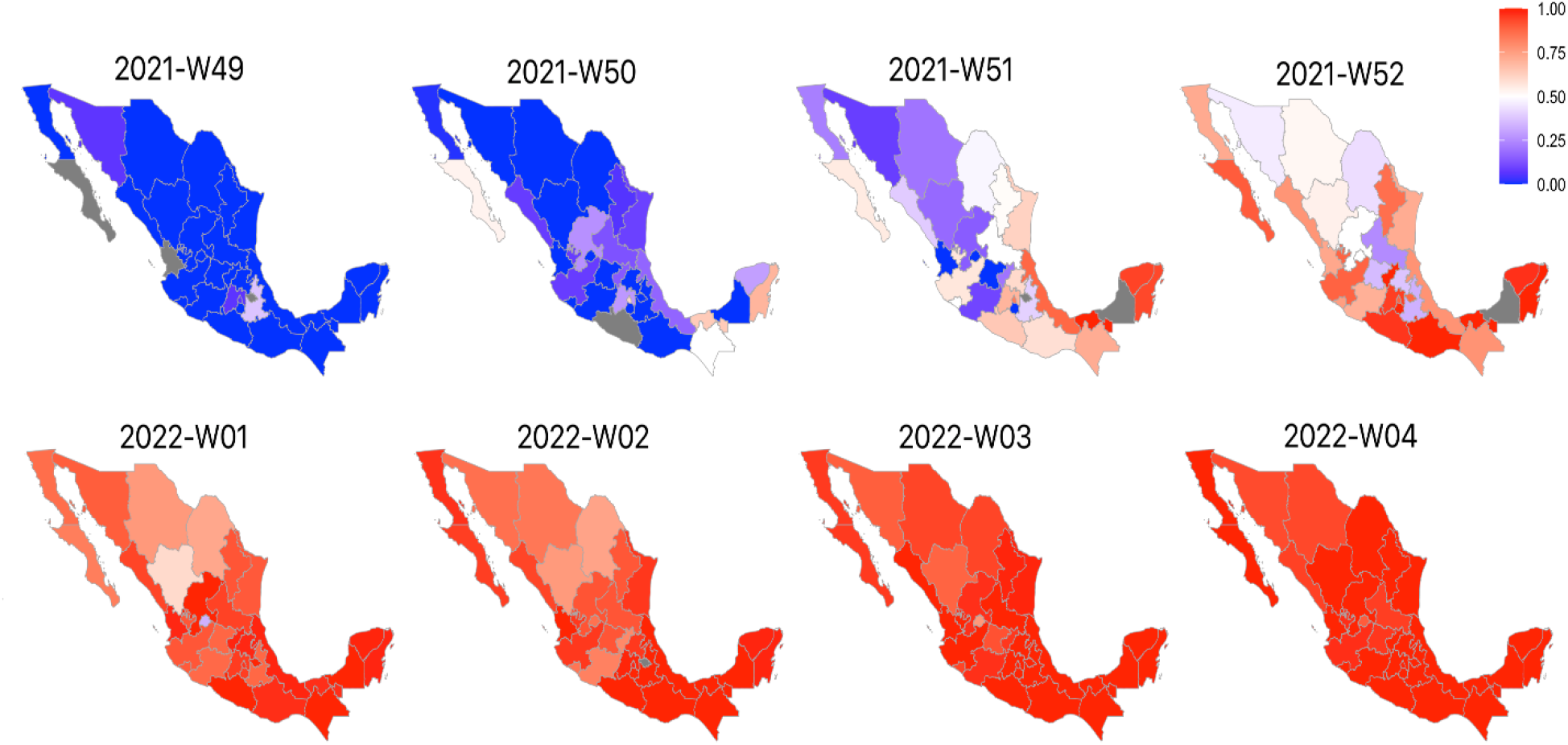
Weekly map series of BA.1.x prevalence in Mexico. According to the legend, the maps show the frequency of BA.1.x sequences per epidemiological week. The red gradient represents the prevalence of Omicron sequences, while the blue gradient represents the prevalence of Delta. States with no available weekly information are shown in gray.

To further characterize the replacement of Delta by BA.1.x throughout the country, we compared four geographical regions, which were determined considering the similarity of weekly lineage abundance in the respective states, as described in the Methods section. The weekly relative frequency of BA.1.x lineages was plotted against the epidemiological week to compare their dynamics of dispersion and prevalence by region (Fig. 3A). Also, for clarity, the number of weeks required to reach a given relative frequency in each area was plotted, beginning with the first week of detection (Fig 3B). A limitation of this analysis was the low number of sequences that were available from the SE region during the initial weeks of the study, so that by the time it was first detected, its prevalence had already ramped up to 50%. To overcome this limitation, the analysis was carried out by comparing the number of weeks it took BA.1.x to increase their prevalence from 50% to 100% in each region. Figure 3B shows that the increase from 50% to 100% required four and six weeks in the SE and CS regions, respectively, but instead took eight weeks in the N and CN regions. In addition to the shorter period it took BA.1.x to dominate the CS and SE, the variant was detected earlier in these regions (Fig 3A).

**Figure 3.**
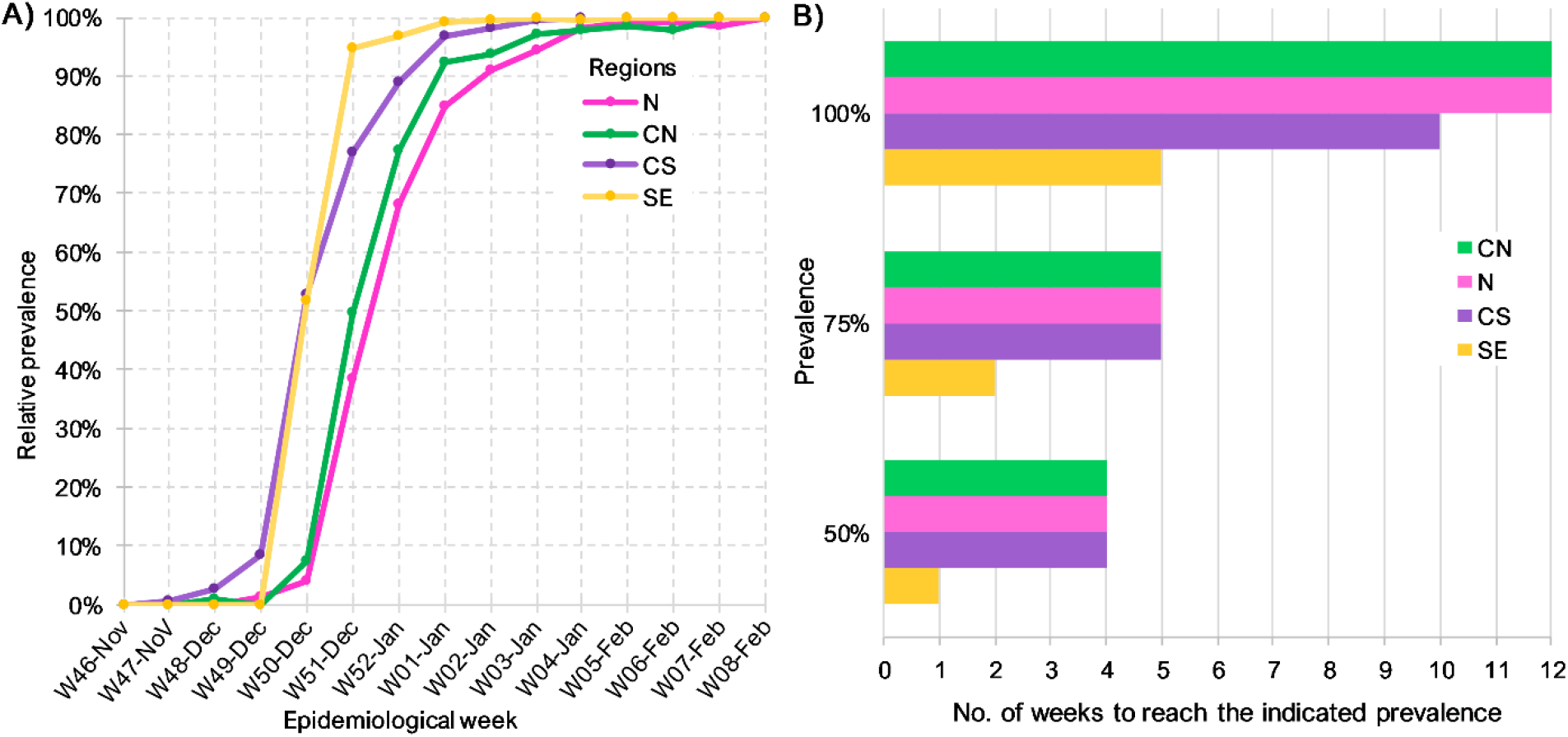
Dynamics of Delta replacement by BA.1.x in different regions of Mexico. A) Prevalence of BA.1.x lineages per epidemiological week. B) Number of weeks since BA.1.x was first detected until it reached the specified prevalence by region.

### Factors affecting BA.1.x dispersion dynamics by region

To understand the differences in the dynamics of BA.1.x dispersion, we analyzed and plotted the sublineage diversity of Delta and BA.1.x per Mexican region from W39-Oct2021 to W05-Feb2022 (Figure 4). Interestingly, the diversity of Delta sublineages in the weeks prior to the entry of BA.1.x also differed by region (W44-Nov to W49-Dec). In the N region, the observed diversity was higher, with six lineages reaching at least 10% of prevalence. In comparison, the CN and SE regions had four and three lineages above 10%, respectively, whereas the CS region exhibited the lowest diversity, clearly dominated by a single sublineage. These results were in agreement with the differences between Shannon diversity indices, which considered the abundance of Delta sublineages from W39-Oct to W48-Nov 2021, to be significantly higher in the N (H=2.3) and CN (H=2.0) regions compared to CS (H=1.7) and SE regions (H=1.8) (N vs. CS, p <0.004; N vs. SE, p = 0.02; CN vs. CS, p<0.01; CN vs. SE, p<0.03).

**Figure 4.**
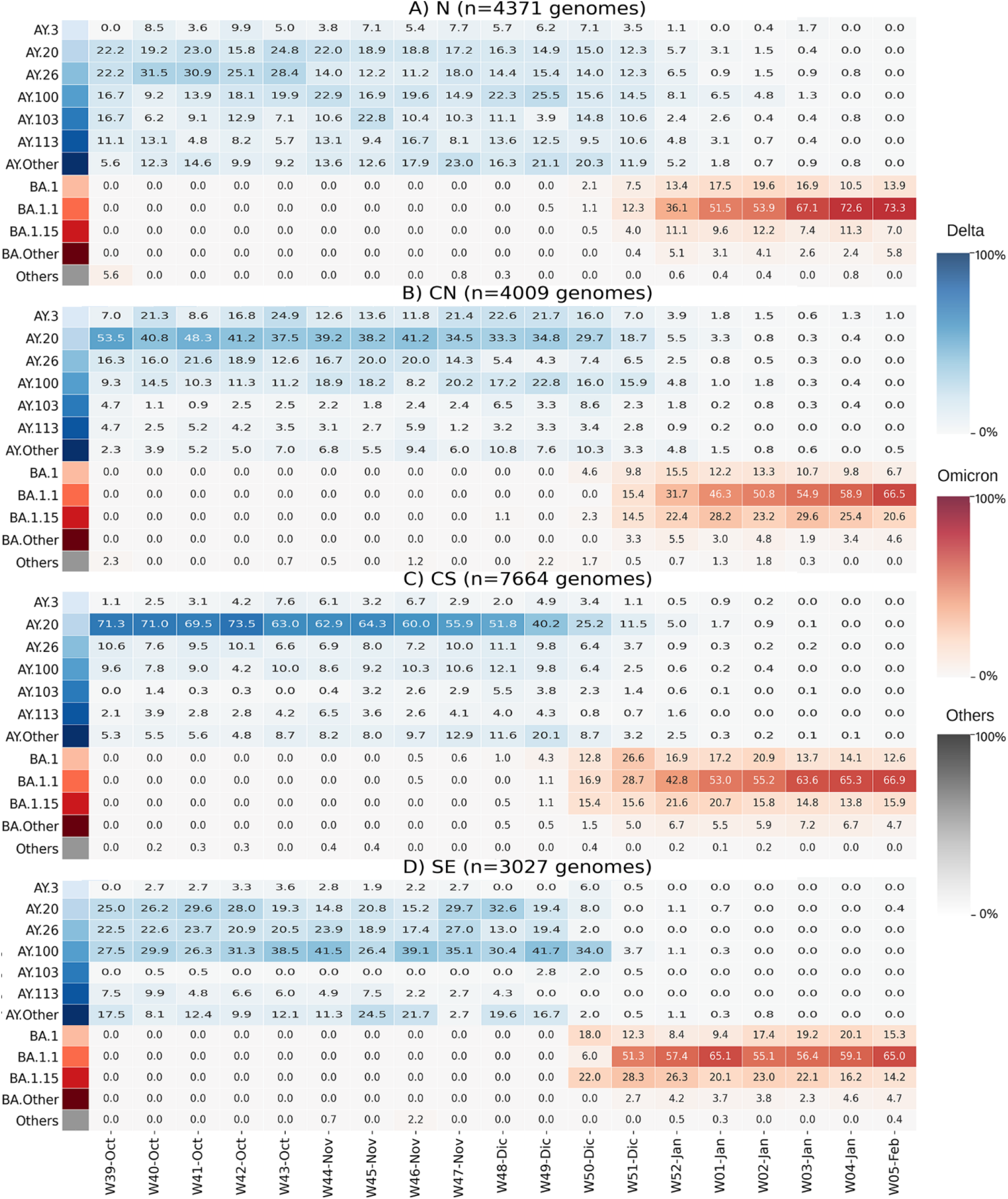
Heatmaps of lineage prevalence per week from October 2021 to February 2022 in the four regions of Mexico: A) North, B) Central North, C) Central South, D) Southeast. Delta sublineages are shown in a blue gradient. Omicron sublineages are shown in a red gradient. All others are collated and shown in gray.

A SARS-CoV-2 high lineage diversity has been associated with population density in Mexico [16]. This factor, rather than lineage diversity *per se*, could explain the differences in BA.1.x dispersion. Accordingly, the N region of Mexico has the lowest population density and the slowest BA.1.x dispersion (see Fig S4). However, the region with the second lowest population density and the fastest spread of BA.1.x was the SE region, suggesting that population density alone cannot determine lineage diversity nor explain disparities in BA.1.x spread rate.

Notably, in the CS and the SE, the proportion of Delta sublineages that circulated during the fourth wave [18], was practically the same than in the inter-wave (W42-Oct to W50-Dec) period. In contrast, in the N region, and to a lesser extent in the CN, the introduction and growth of additional Delta sublineages occurred during the inter-wave interval, such as AY.3, AY.103, and AY.113 (Fig 4).

To investigate other factors contributing to the variations in the BA.1.x spread, aside from the diversity of Delta lineages before the onset of BA.1.x and population density in the regions, the epidemic curves of these regions were examined (Figure 5). The epidemic curve of the N region shows an active inter-wave interval (Fig 5A). The maximum number of cases in N during the Delta wave was 1,880 per day, and during the inter-wave period, cases persisted at an average of 977 per day, corresponding to 52% of the Delta wave’s peak. Additionally, infections in this region began to increase during epidemiological W47-Nov and peaked at 1,170 daily cases during W48-Nov. Throughout this period, all sequences identified in the N region corresponded to the Delta variant (see Fig 4A). In contrast, in the CN region, SARS-CoV-2 circulation during the Delta-BA.1.x inter-wave period was roughly 13% of the Delta peak, whereas it was 10% and 6% in the CS and SE regions, respectively (Figure 5B, C, and D).

**Figure 5.**
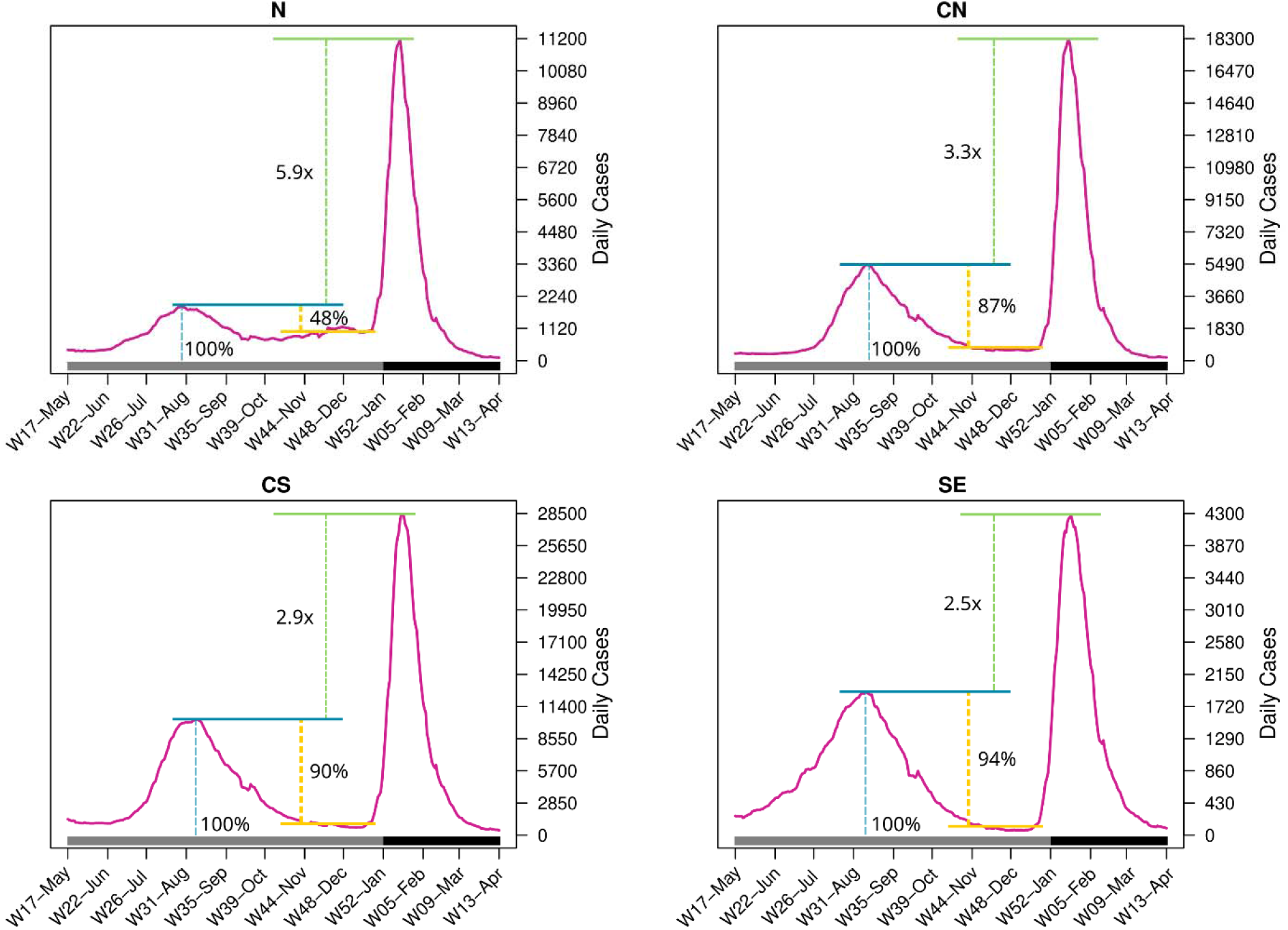
Number of daily confirmed cases in the four regions per epidemiological week. The bottom horizontal bar marks the year (gray, 2021; dark gray, 2022).

Additionally, in the N region, the BA.1.x peak was 5.9 times higher than Delta’s, while in the CN region, it was 3.3 times higher. The increase in the CS and SE regions was 2.9 and 2.5, respectively, resulting in a smaller disparity. Taking all data together, a higher Delta peak (relative to the BA.1.x peak), lesser virus circulation during the inter-wave period, and/or a lower Delta variant diversity were associated with a faster replacement of Delta by BA.1.

## Discussion

In this study, we analyzed the SARS CoV2 genome sequences isolated in Mexico around the time of the introduction of the Omicron-BA.1 variant to the country to better understand its spread during the pandemic’s fourth wave in Mexico. The first national case was reported by late November 2021, quickly becoming the dominant variant by mid-December. The first significant observation was the rapid increase in the number of infected people, accounting for the highest peak thus far in the pandemic, occurring in mid-January 2022 with over 62,000 daily infections. In contrast, and as was reported in other countries, this variant had a relatively minor impact on hospitalizations and deaths. This phenomenon has been attributed to high vaccination rates and a general reduction in the susceptible population, despite the ability of the BA.1 virus to evade neutralization, and to Omicron’s potentially lower virulence [2,28–30].

The spread of BA.1.x sublineages in Mexico did not result in the acquisition of specific new mutations nor the clustering of sequences by sublineage or geographical region, in contrast to previous reports on the spread of Alpha and Delta VOCs, which described the emergence of local specific mutations and sublineages, as well as a solid geographical association with those changes [17,18]. The low diversity of Omicron sublineages was probably associated with its unprecedentedly rapid spread across the country. This observation is consistent with a previous report describing BA.1.x entry and spread in Mexico City [31], and with descriptions from other countries. For instance, in an analysis of South American Omicron sequences, the authors divided them into clusters using phylogenetic methods; however, these clusters contained sequences from different sublineages and countries, suggesting an overall low diversity [32]. Additionally, in Finland, a phylogenetic analysis of BA.1.x sequences reported multiple singletons and low clustering of Finnish sequences [33].

Despite the reported transmission advantage of BA.1 over Delta [1,2,5], this variant propagated more slowly in the N and CN of Mexico than in the CS and SE regions. The slower spread rate may be attributed to a higher diversity of Delta sublineages and epidemic activity during the BA.1.x inter-wave interval. Notably, in the N region, the number of active SARS-CoV-2 cases reported during the Delta-BA.1.x inter-wave period was the highest in the country, representing 52% of the maximum peak of the Delta wave in that region, indicating continued transmission during the fall of 2021. The active circulation of the virus may have allowed the introduction of new Delta sublineages, sustained by the low population density and a low proportion of people with prior Delta infection. Consequently, this region exhibited the highest diversity of Delta sublineages before the onset of Omicron.

Conversely, CS and SE had the lowest number of cases in their Delta-BA.1.x inter-wave period and the smallest differences in the ratio between the peaks of the third and fourth waves. Additionally, these regions had the lowest diversity of Delta lineages at the onset of Omicron. Also, in contrast to the N region, no change was observed between the sublineages circulating previously during the Delta peak and in the inter-wave period [18]. The CN region seems to be a transition zone between the N and the CS and SE regions, as it shares a slower replacement and a higher diversity of Delta with the N region. Still, its epidemic curve was comparable to that of the CS and SE regions. The association between the epidemiological and genomic profile of the viruses in a region with the rate of Omicron dispersion has not been reported to our knowledge.

Geographical and prevalence differences in the circulation of distinct lineages in Mexico have been previously reported [16–18]. The prevalence of B.1.1.519, which dominated the center and south of the country between February and May 2021, was lower in the north [17]. Conversely, in the north of Mexico, the Alpha variant circulated at a higher prevalence than in the rest of the country during roughly the same period. Whereas the Alpha variant had a proven transmission advantage and managed to displace other lineages in many countries, the B.1.1.519 lineage only was dominant in Mexico, suggesting that its fitness advantage was less than that of Alpha. This observation indicates that fitness alone is insufficient to predict the spread of a SARS-CoV-2 variant, which is, in fact, a complex phenomenon. In this study, Delta lineage diversity seems to have influenced the rate of spread of Omicron-BA.1.x in Mexico. However, the sublineages that circulated in these regions could also play an important role in this regionally delayed replacement. Sublineages AY.3, AY.103, and AY.113 circulated at higher frequencies in the N and CN, which may have interfered with the arrival of BA.1.x. Particularly, AY.3 and AY.103 have been associated with breakthrough infections in vaccinated people, with AY.3 having caused asymptomatic cases with high viral loads [34] and AY.103 having a breakthrough rate of 41%, higher than other Delta sublineages [35]. Therefore, these viruses could continue circulating during the inter-wave period and possibly delay the entry of BA.1. Of note, a molecular dynamic simulation has shown that AY.3 may exhibit a higher affinity for ACE2 than Omicron-BA.1 [36], further suggesting that some Delta sublineages may have higher fitness and, thus, be maintained in the population longer.

The importance of Delta infections during the inter-wave period on the rate of spread of Omicron-BA.1.x remains unclear, even though the level of Delta-Omicron inter-wave virus circulation, as compared to the preceding Delta-peak in the different regions of the country was inversely correlated with the spread of the BA.1.x variants.

## Author contributions

## Acknowledgments

We are grateful to the authors from the originating laboratories responsible for obtaining the samples and the submitting laboratories responsible for generating the genomes shared via GISAID, on which this research is based. For a full list of contributors of each sequence and its details including the Originating Lab and Submitting Lab and the list of Authors, visit https://doi.org/10.55876/gis8.220927gw. We appreciate the computer assistance provided by Jerome Verleyen, Roberto Bahena, Juan Manuel Hurtado, and Verónica Jacinto. We thank Julissa Enciso-Ibarra and Karla G. Aguilar-Rendon for technical help. The project LANCAD-UNAM-DGTIC-350 of the Dirección General de Cómputo y Tecnologías de la Información (DGTIC-UNAM) provided Supercomputing resources in Miztli.

