## SupplementalFigures for "Regional epidemic dynamics and Delta variant diversity resulted in varying rates of spread of Omicron-BA.1 in Mexico"

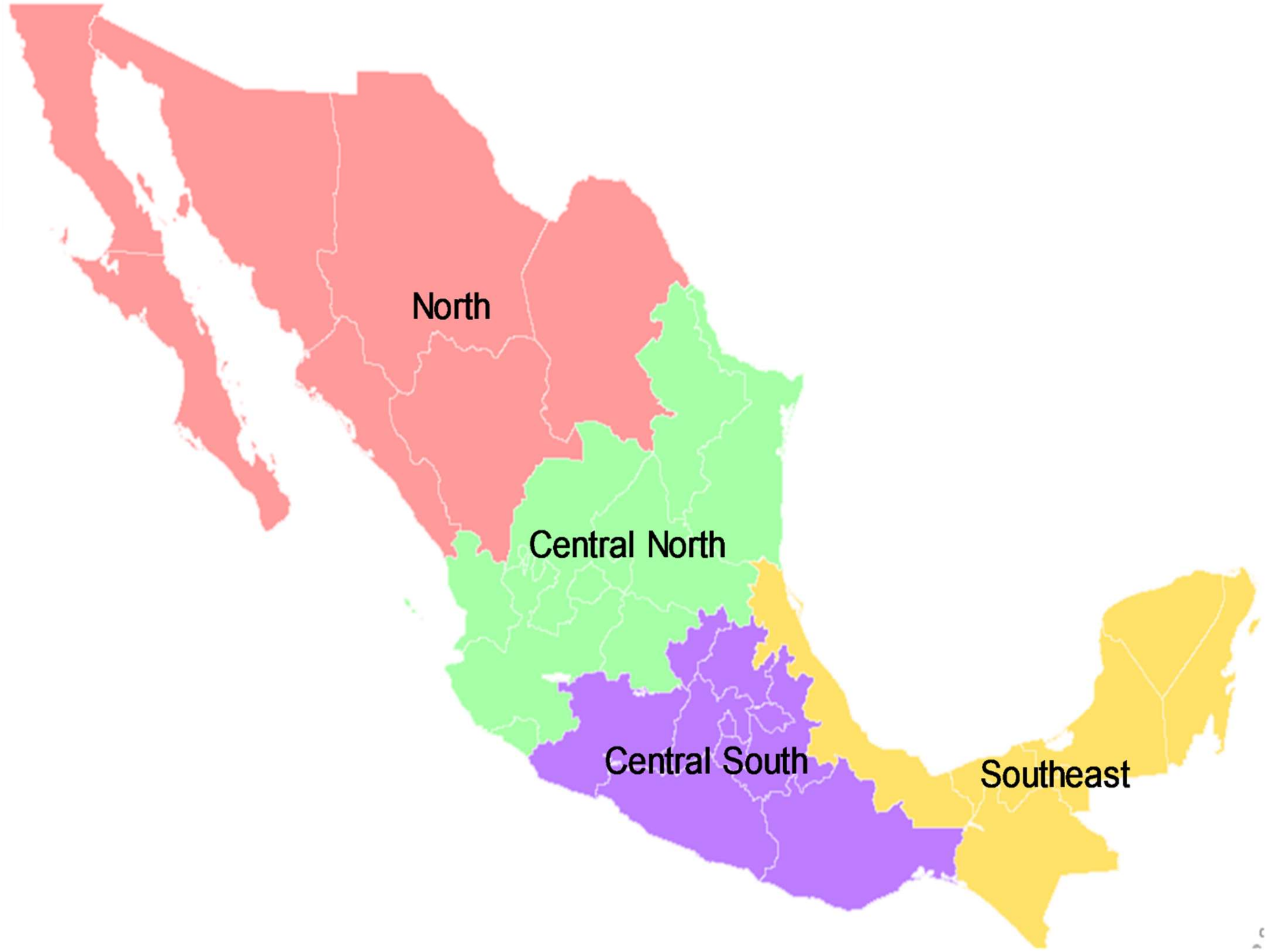


**Figure S1.** Map of the 32 States in Mexico with the four geographical regions compared in the study.

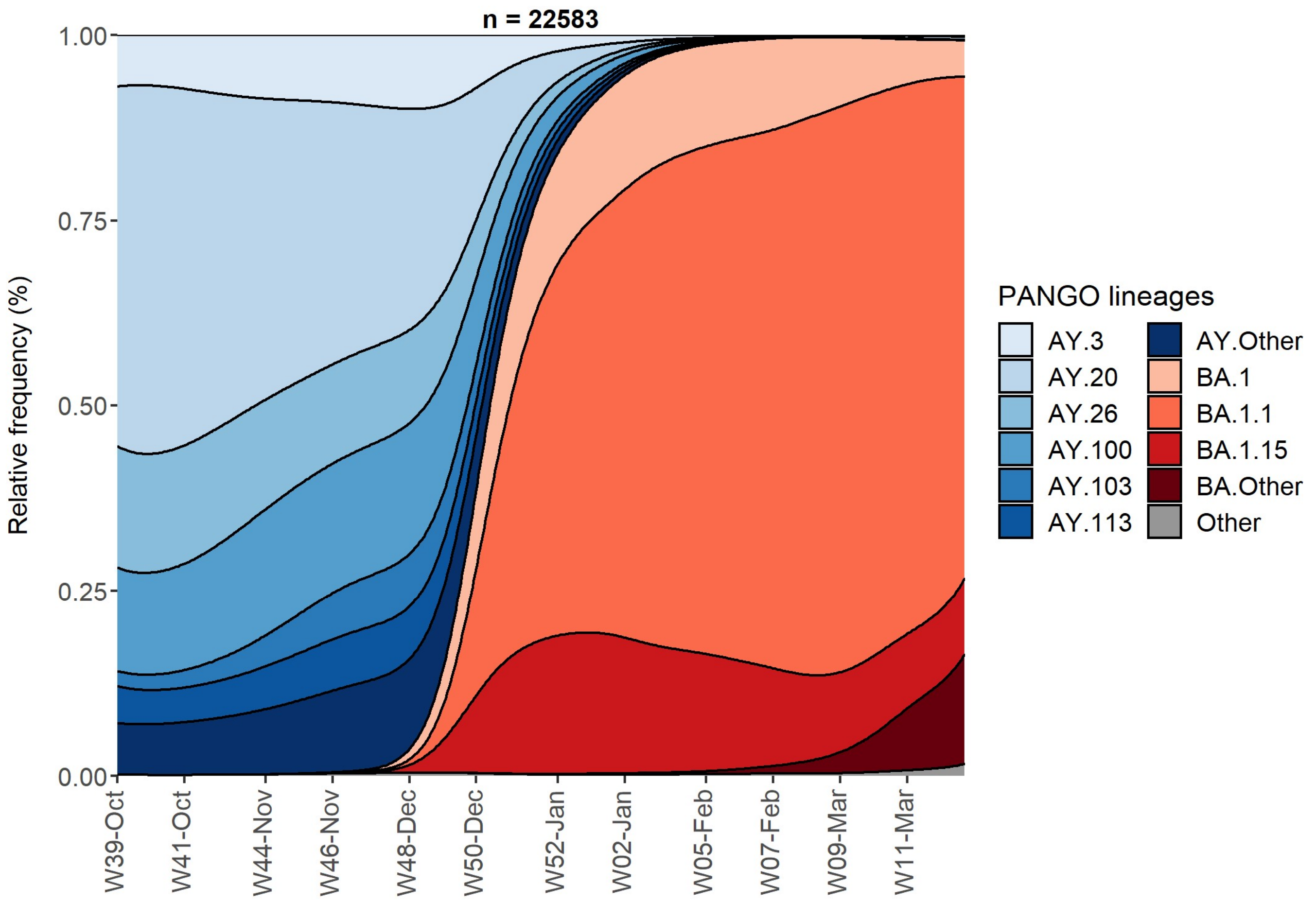


**Figure S2**. Stacked density plot of the weekly relative frequency of SARS-CoV-2 lineage circulation in Mexico from October 2021 until March 2022

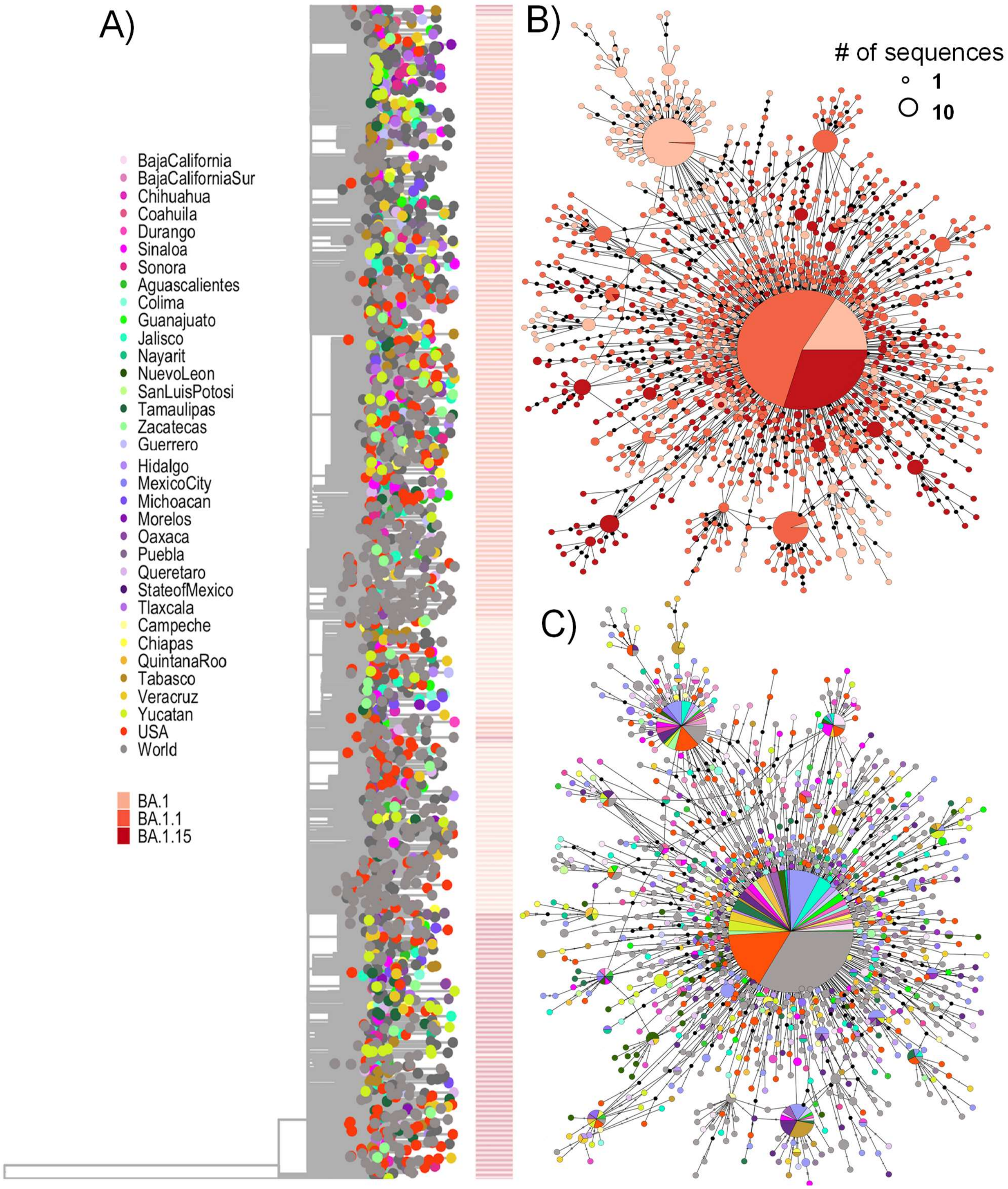


**Figure S3.** Phylogeny and haplotype network of BA.1, BA.1.1, and BA.1.15 Omicron sublineages. Sequences from these sublineages from Mexico and abroad were used. A) Time-scaled maximum-likelihood phylogeny. The tips are colored by the state of sampling and a heatmap with the sublineage is shown on the side. B and C) Haplotype network colored by sublineage and state of sampling, respectively. The size of the circles corresponds to the number of samples within the same haplotype (scale is provided). The black circles in the haplotype networks represent mutations between sequences.

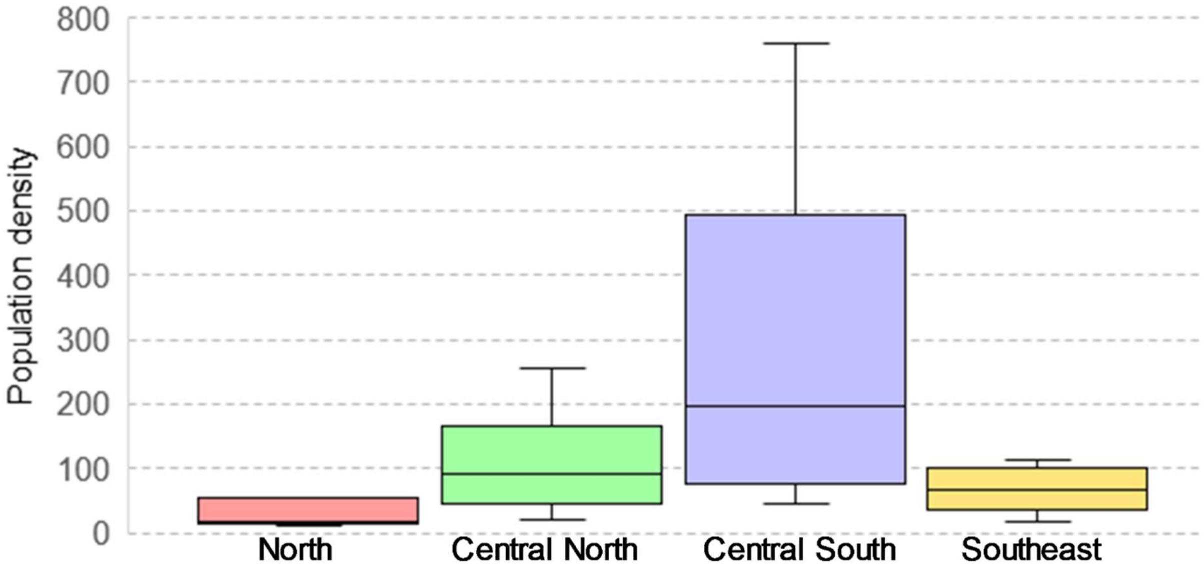


**Figure S4.** Distributions of the population density in the four regions in Mexico. The density was calculated as inhabitants per square kilometer (0.62 miles). Each box shows the interquartile range (IQR) and the median, whilst the whiskers depend on the kurtosis. Mexico City, with density over 6000, was considered an outlier (further than 1.5 times the IQR from the boxes) and is not shown.
